# Barriers to Integration of Bioinformatics into Undergraduate Life Sciences Education

**DOI:** 10.1101/204420

**Authors:** Jason J. Williams, Jennifer C. Drew, Sebastian Galindo-Gonzalez, Srebrenka Robic, Elizabeth Dinsdale, William Morgan, Eric W. Triplett, James Burnette, Samuel Donovan, Sarah Elgin, Edison R. Fowlks, Anya L. Goodman, Nealy F. Grandgenett, Carlos Goller, Charles Hauser, John R. Jungck, Jeffrey D. Newman, William Pearson, Elizabeth Ryder, Melissa A. Wilson Sayres, Michael Sierk, Todd Smith, Rafael Tosado-Acevedo, William Tapprich, Tammy C. Tobin, Arlín Toro, Lonnie Welch, Robin Wright, David Ebenbach, Mindy McWilliams, Anne G. Rosenwald, Mark A. Pauley

## Abstract

Bioinformatics, a discipline that combines aspects of biology, statistics, and computer science, is increasingly important for biological research. However, bioinformatics instruction is rarely integrated into life sciences curricula at the undergraduate level. To understand why, the Network for Integrating Bioinformatics into Life Sciences Education (NIBLSE, “nibbles”) recently undertook an extensive survey of life sciences faculty in the United States. The survey responses to open-ended questions about barriers to integration were subjected to keyword analysis. The barrier most frequently reported by the ~1,260 respondents was lack of faculty training. Faculty at associate’s-granting institutions report the least training in bioinformatics and the least integration of bioinformatics into their teaching. Faculty from underrepresented minority groups (URMs) in STEM reported training barriers at a higher rate than others, although the number of URM respondents was small. Interestingly, the cohort of faculty with the most recently awarded PhD degrees reported the most training but were teaching bioinformatics at a lower rate than faculty who earned their degrees in previous decades. Other barriers reported included lack of student interest in bioinformatics; lack of student preparation in mathematics, statistics, and computer science; already overly full curricula; and limited access to resources, including hardware, software, and vetted teaching materials. The results of the survey, the largest to date on bioinformatics education, will guide efforts to further integrate bioinformatics instruction into undergraduate life sciences education.

## Significance Statement

Bioinformatics, a discipline combining biology, statistics, and computer science, is increasingly important for biological research. However, few students in the life sciences learn bioinformatics as undergraduates. Data collected from a survey of life sciences faculty in the United States (*n* = 1,264), the first large-scale study of its kind, demonstrate that there are significant barriers to integration of bioinformatics instruction. The most frequently reported barrier was limited faculty training. Other issues included lack of student interest in bioinformatics; lack of student preparation in mathematics, statistics, and computer science; an overly full curriculum; and limited access to resources, including hardware, software, and vetted teaching materials. The information from this survey can guide efforts to increase integration of bioinformatics into life sciences education.

## Introduction

Bioinformatics, a discipline combining aspects of biology, statistics, and computer science, is becoming increasingly important for research efforts in all areas of biology (1, 2). Students graduating with some experience in bioinformatics have more employment opportunities available to them (3) and are better prepared for graduate studies in life science fields. It has also been suggested that students graduating with degrees in molecular biology and biochemistry should have some familiarity with bioinformatics (4). With the growing emphasis on “big data” in biology, there is more demand for researchers in the life sciences with training in bioinformatics. Yet many life science students earn their degrees without any exposure to bioinformatics (5, 6).

The Network for Integrating Bioinformatics into Life Sciences Education (NIBLSE, “nibbles”) is a group of US educators in biology, bioinformatics, and computer science, as well as professionals from the private sector, dedicated to making bioinformatics an integral component of instruction in the life sciences nationwide. NIBLSE is supported by an Undergraduate Biology Education Research Coordination Network grant from the National Science Foundation (NSF) and aims to develop instructional strategies for undergraduates to gain experience in bioinformatics. At the same time, NIBLSE is working to address barriers to the implementation of those strategies (7). The Netherlands-based Global Organisation for Bioinformatics Learning, Education, and Training (GOBLET), is also examining the bioinformatics education landscape (8) but has not specifically addressed barriers to bioinformatics training at the undergraduate level. In the United States, bioinformatics instruction has predominately been provided at the graduate level and, in some instances, as a stand-alone course at the undergraduate level (9–11). However, there has been little effort to integrate bioinformatics broadly into the undergraduate curriculum.

To further the integration of bioinformatics into life sciences education, we need to understand the barriers preventing broad implementation of bioinformatics. To date, there have been no large-scale studies examining faculty views on bioinformatics in undergraduate biology curricula, particularly in the context of the needs of different types of institutions. To that end, NIBLSE surveyed biology faculty at two-year and four-year institutions primarily in the United States, gathering more than 1,260 responses. The survey included several open-ended, free-response questions that focused on barriers to the integration of bioinformatics instruction. Responses to these questions were subjected to keyword analysis. The results provide a national view on several key barriers. We discuss each of the major barriers uncovered and describe efforts by NIBLSE to alleviate these barriers.

## Results

NIBLSE is dedicated to increasing the integration of bioinformatics into life sciences education. We therefore sought to determine the extent to which the US life sciences community shares the view that bioinformatics is essential to life sciences education. We received more than 1,260 responses to our survey, the vast majority from the United States; demographic information is shown in Fig. 1. Approximately 95% agreed with the statement that “bioinformatics should be integrated into undergraduate life sciences education.” At the same time, 60% of respondents said that they do not currently teach courses with substantial bioinformatics content.

**Fig. 1.**
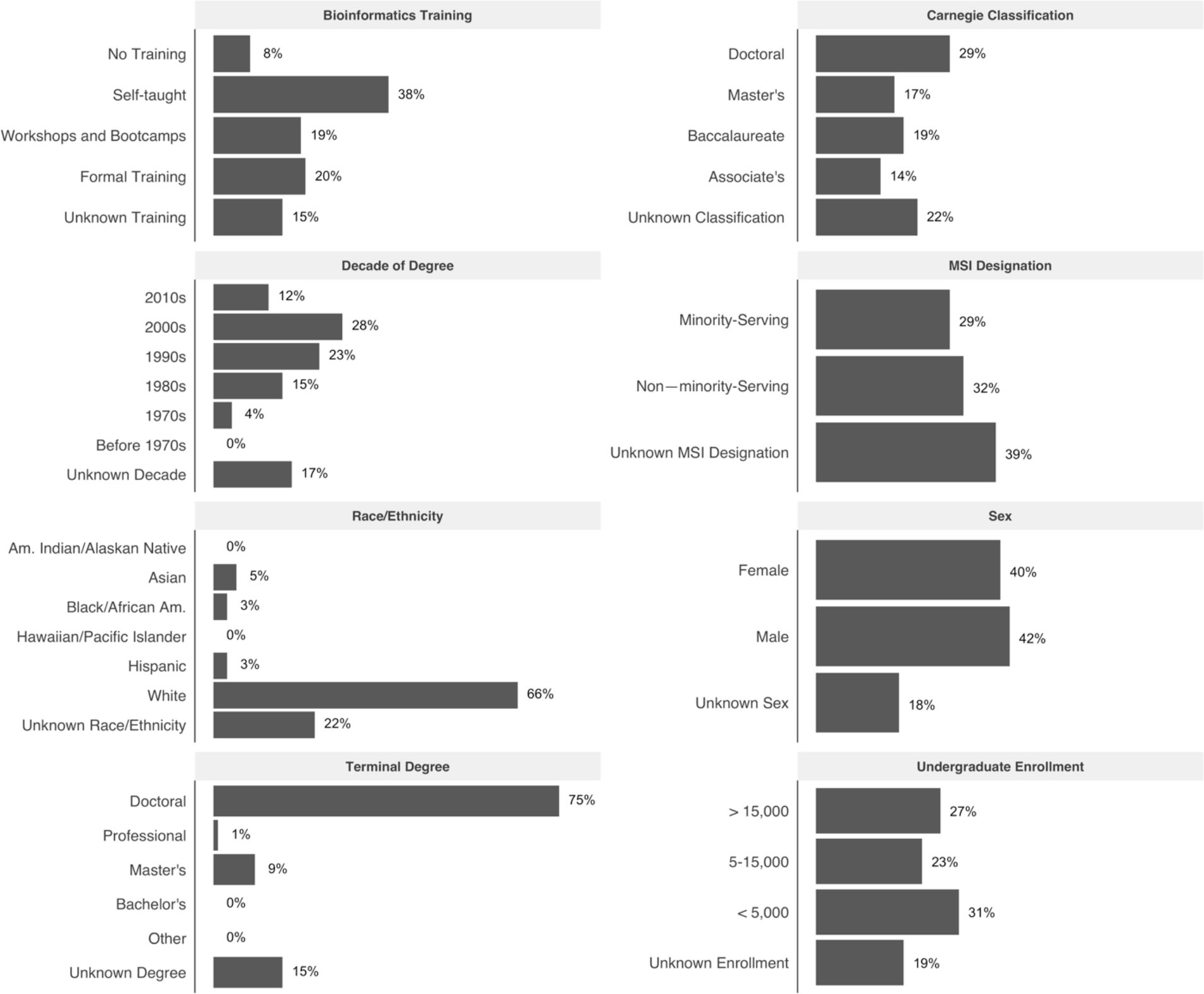
Summary Demographics. Summary demographics shown as percentages of respondents. The composite survey respondent is a white male or female PhD, self-taught in bioinformatics, with a PhD degree earned in 2000-2009. S/he works at a non-minority-serving, doctoral-granting institution with an undergraduate enrollment of less than 5,000. In subsequent analyses, unknown or undisclosed values were not considered. (*n* = 1,231, the total US respondents).

Respondents had the opportunity to provide comments to four free-response questions about barriers to including bioinformatics in their teaching (Table 1). As described in Materials and Methods, these qualitative responses were coded according to keywords and then binned according to super-category (for example, “Faculty Issues”) and category within that supercategory (for example, within “Faculty Issues,” one of the bins was “Lack of training”). Three of the four questions (Questions 1, 2, and 3) prompted responses that fell into the same supercategories. (Super-categories and categories for Question 1 are listed in Table S1; all analyses are available at GitHub: https://github.com/niblse/barriers_to_bioinformatics_integration) Question 4, which focused specifically on technical barriers, elicited a different set of responses and was coded differently. Although not every respondent answered any or all of these open-ended questions, we coded nearly 2,000 responses for these four questions (Table S2). In the sections below, we describe the findings from the survey with respect to the super-categories “Faculty Issues” and “Student Issues,” then describe some additional results with respect to other barriers frequently mentioned by the respondents.

**Table 1.**
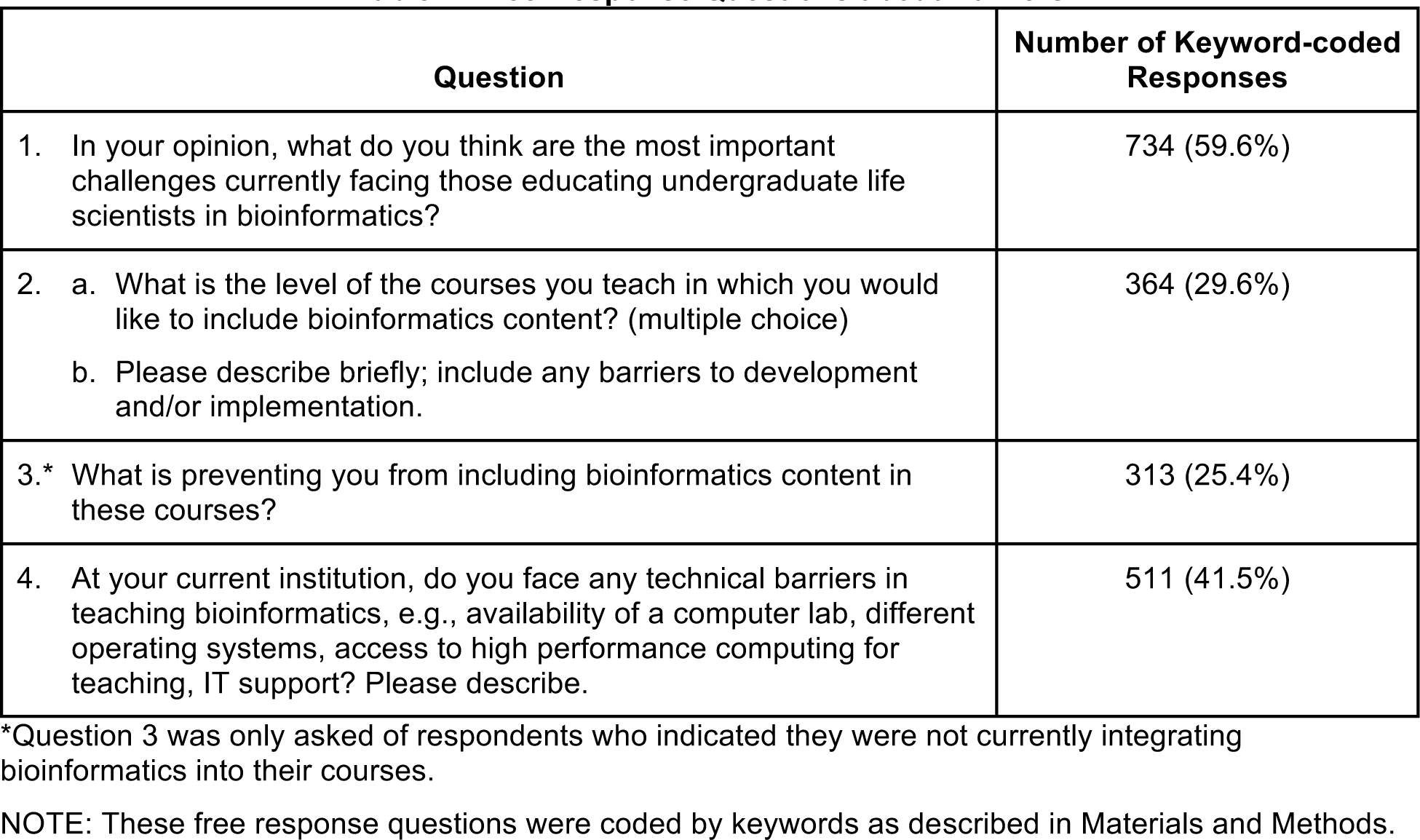
Free-Response Questions about Barriers.

| Question | Number of Keyword-coded Responses |
| --- | --- |
| 1. In your opinion, what do you think are the most important challenges currently facing those educating undergraduate life scientists in bioinformatics? | 734 (59.6%) |
| 2. a. What is the level of the courses you teach in which you would like to include bioinformatics content? (multiple choice)<br>b. Please describe briefly; include any barriers to development and/or implementation. | 364 (29.6%) |
| 3.* What is preventing you from including bioinformatics content in these courses? | 313 (25.4%) |
| 4. At your current institution, do you face any technical barriers in teaching bioinformatics, e.g., availability of a computer lab, different operating systems, access to high performance computing for teaching, IT support? Please describe. | 511 (41.5%) |

\*Question 3 was only asked of respondents who indicated they were not currently integrating bioinformatics into their courses.
NOTE: These free response questions were coded by keywords as described in Materials and Methods.

### Super-Category: Faculty Issues

As shown in Fig. 2 and Fig. S1, the lack of faculty training topped the list of barriers. Many faculty also mentioned the lack of instructional time as a significant barrier. Although lack of faculty training was the most significant barrier at most institution types, as noted in Fig. 3, faculty at doctoral-granting institutions reported this barrier less frequently than those at other institution types. Faculty at doctoral-granting institutions also rated advanced technical skills, such as those in statistics and scripting, to be more important competencies than did faculty at other types of institutions (submitted, accessible at http://www.biorxiv.org/content/early/2017/08/03/170993).

**Fig. 2.**
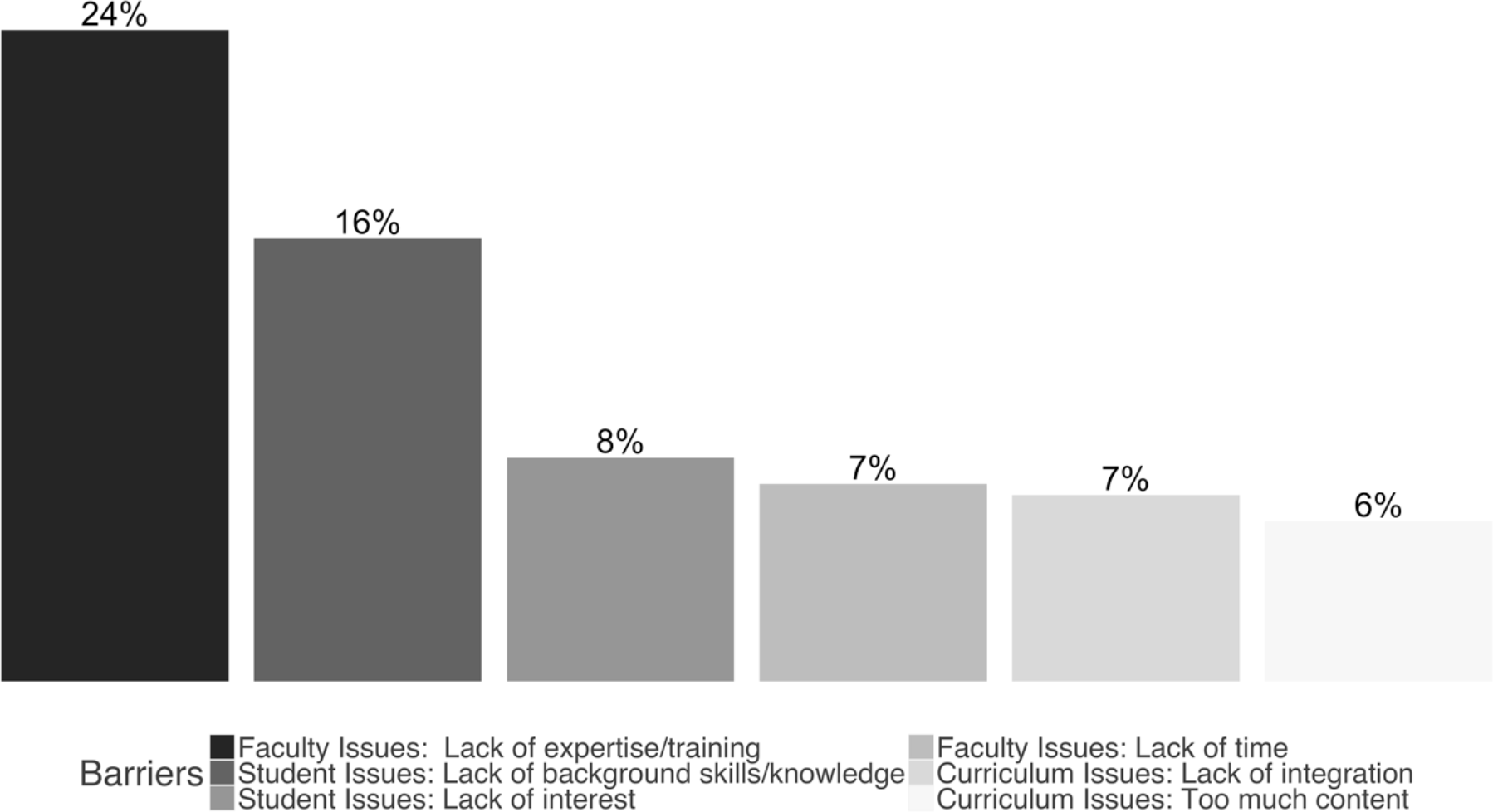
Summary of Most Commonly Reported Barriers by Category. The most commonly reported barriers were lack of faculty expertise and training (24%), followed by students’ lack of background skills and knowledge (16%), student lack of interest (8%), faculty lack of time (7%), incompatibility of bioinformatics with current curriculum (7%), and too much existing content (6%). (*n* = 1,231; 734 coded responses).

**Fig. 3.**
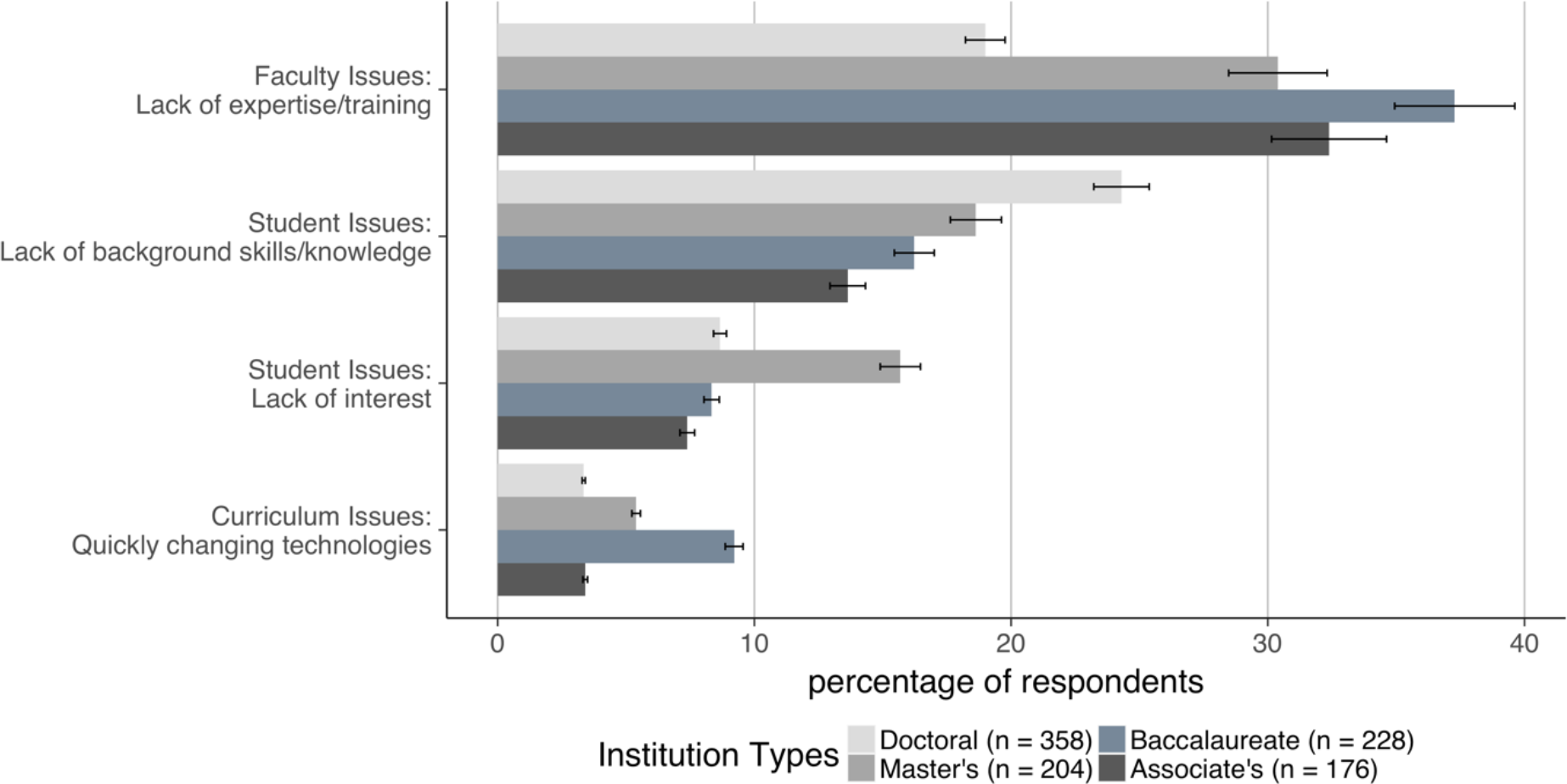
Differences in Barrier Perceptions by Institutional Carnegie Classification. Four barriers were reported differently by faculty at the different institution types. Faculty Expertise and Training (*P* = 8.25e-06) was mentioned most frequently by faculty at baccalaureate-granting institutions (37% vs. 19% at doctoral-granting institutions). Student lack of background skills and knowledge (*P* = 0.01) was mentioned most frequently by faculty at doctoral-granting institutions (24% vs.14% at associate’s-granting institutions). Student lack of interest (*P* = 0.02) was mentioned most frequently by faculty at master’s-granting institutions (16% vs. 7% at associate’s-granting institutions). Curriculum issues caused by quickly changing technologies (*P* = 0.01) was mentioned most frequently by faculty at baccalaureate-granting institutions (9% vs. 3% at associate’s-granting institutions). (*n* = 966, effect size at 80% = 0.106, meaning medium effects detected).

To illustrate these barriers—lack of faculty training and lack of instructional time—we include relevant quotes from survey respondents below. (The comments have been lightly edited for clarity and to correct typographical errors.)

- *Faculty issues: Lack of expertise/training* “Lack of training in the area, as many of us earned our PhDs before bioinformatics was widely used and available.” “Lack of faculty expertise in the rapidly changing disciplines of genomics/bioinformatics and related technologies.” “A lack of training and experience in bioinformatics.”
- *Faculty issues: Lack of time* “A lack of agreement about what topics should be dropped in order to have space/time to introduce bioinformatics.” “Time to fit into the curriculum.” “Too much other stuff in the Biology curriculum that is perceived as essential.”

We hypothesized that faculty who earned their highest degree most recently would report the most formal training in bioinformatics. This hypothesis was confirmed: 48% of faculty who earned their degree in 2010-2016 reported they had some kind of formal training (undergraduate or graduate courses and/or certificates) compared to 8% of faculty from the 1980-1989 cohort (Table 2). Despite this level of formal training, faculty who earned degrees in 2010-2016 were the least likely (*P* = 0.003) to report teaching dedicated bioinformatics courses or integrating bioinformatics into their teaching. (Only 25% of the 2010-2016 cohort report integrating bioinformatics into their classrooms compared to 42% of respondents in the 1990–1999 and 2000-2009 cohorts or 35% of respondents in the 1980-1989 cohort.) Although faculty from the 2010-2016 cohort teach at all institution types in about the same percentages (Table S3), it may be that as new faculty, they are unable to shape the overall curriculum and/or are not tasked with teaching courses that match their skills in bioinformatics (Fig. 4).

**Table 2.**
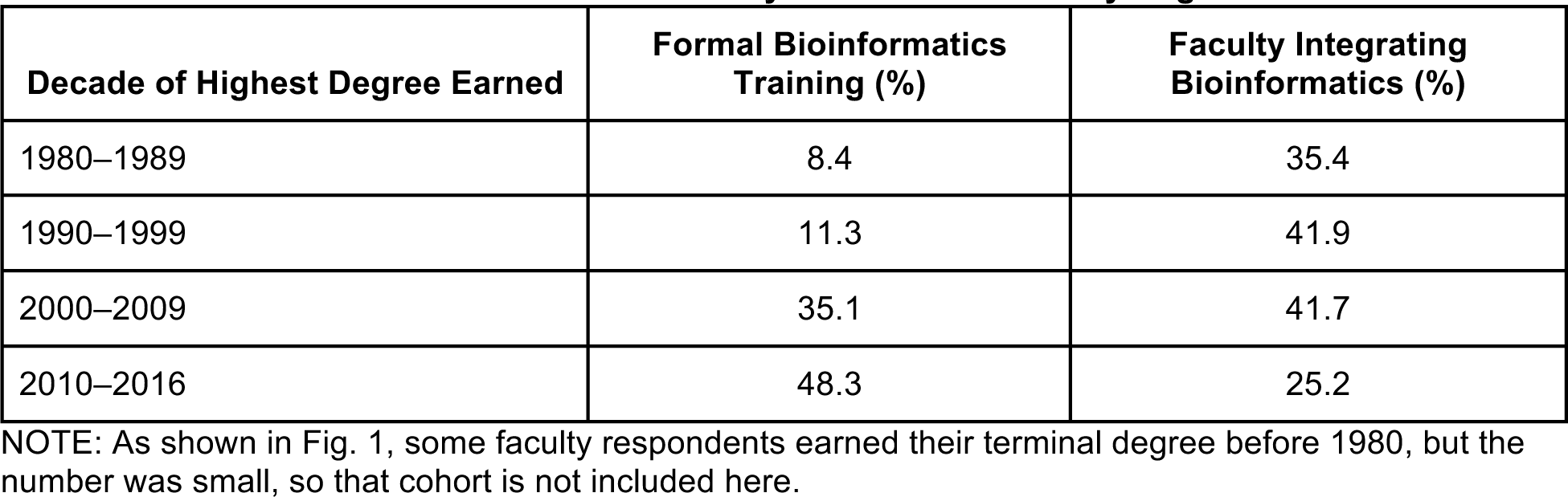
Characteristics of Faculty Cohorts Stratified by Degree Year.

**Fig. 4.**
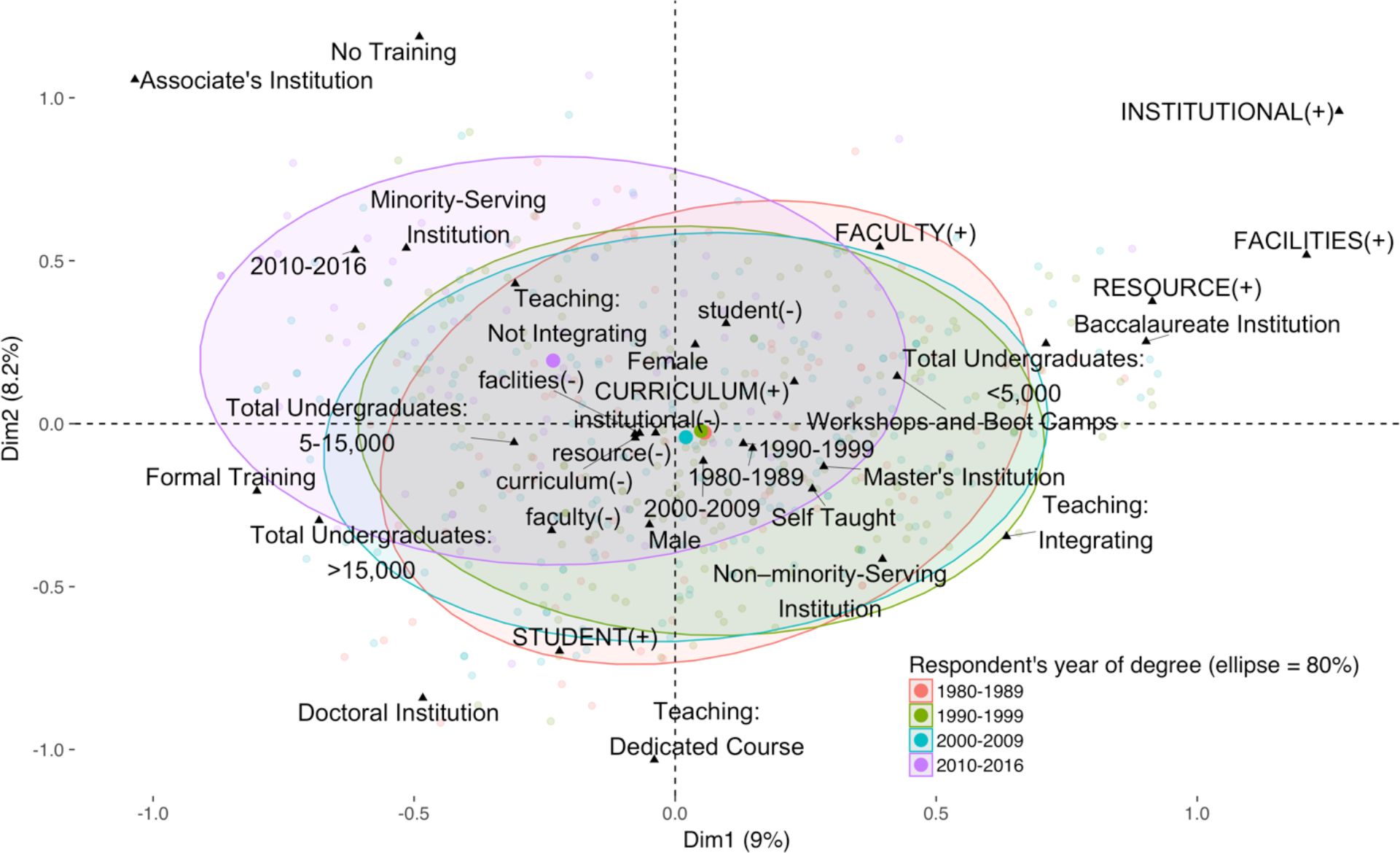
Barriers Reported by Faculty Members Stratified by Year of Highest Degree. Multiple correspondence analysis (an analysis similar to principle component analysis—see Materials and Methods) was calculated for several demographic characteristics. Here, respondents are grouped by decade of terminal degree in a colored ellipse encompassing 80% of individuals in a sub-demographic (the center of which is marked by a large colored dot). The youngest cohort (2010-2016) separates from the older cohorts. Several qualitative characteristics (black triangles) are mapped, and their proximity to each other is an indication of their tendency to be reported together (e.g., non-minority-serving institution and integrating bioinformatics tend to be reported together, as seen in the lower-right quadrant, and minority-serving institution and not integrating bioinformatics tend to be reported together, as shown in the upper-left quadrant). Here we map: 1) level of bioinformatics training (No Training, SelfTaught, Workshops and Boot Camps, Formal Training); 2) current bioinformatics content in teaching (Teaching: Dedicated Bioinformatics Course, Teaching: Integrating Bioinformatics, Teaching: Not Integrating Bioinformatics); 3) sex (Female, Male); 4) institution minority-serving status (Minority-serving Institution, Non-Minority-Serving Institution); 5) undergraduate enrollment (Total Undergraduates < 5,000, Total Undergraduates 5-15,000, Total Undergraduates > 15,000). Finally, we display responses to Barriers Question 1 as qualitative supplementary values (identified as ‘CATEGORY (+)’ when an issue was reported in that supercategory or as ‘category (−)’ when no issue was reported; e.g., FACULTY (+) indicates a faculty-related barrier). (*n* = 526).

We also examined the data with respect to whether the respondent was a member of an underrepresented minority in STEM as defined by the National Science Board (12), but it should be noted that the number of respondents identifying as African American, Hispanic, American Indian/Alaskan Native, or Native Hawaiian or other Pacific Islander made up less than 7% of our sample (81 respondents), mirroring the lack of diversity in life science faculty in the United States (13). We grouped underrepresented minorities into a single group sufficient for analysis. Members of underrepresented groups report training as a barrier more frequently than members of non-underrepresented groups, with 41% of STEM-minority faculty reporting a training barrier vs. 28% of faculty not underrepresented in STEM (*P* = 0.01, *n* = 961, effect size at 80%, power = 0.09, meaning small effects were detected). This result is consistent with our findings that faculty from underrepresented groups rated every competency as more important than faculty from groups that are not underrepresented in STEM (submitted, accessible at http://www.biorxiv.org/content/early/2017/08/03/170993).

Comparing faculty at minority-serving institutions (MSIs) with those at non-MSIs, MSI faculty report faculty issues as a barrier at a slightly, but not significantly, lower rate than faculty at non-MSIs (Fig. S2). This may be explained by the lower number of faculty at MSIs who are integrating bioinformatics; only 23% of the faculty at MSIs are integrating bioinformatics into their teaching in some way compared to 43% of faculty at non-MSIs. With respect to sex, females report training as a barrier slightly more than males do (16% vs. 11%), and they also report lack of access to computer labs at double the percentage of males reporting this problem (Question 4, Table 1; Fig. S3).

### Super Category: Student Issues

The super-category “Student Issues” was the second most frequently mentioned set of barriers after Faculty Issues (Fig. 2). Two particular issues were noted. First, students’ lack of background skills and knowledge, specifically in computer science and statistics, and second, students’ lack of interest in the topic. Faculty at doctoral-granting institutions cited students’ lack of knowledge at a higher rate while faculty at master’s-granting institutions more frequently cited students’ lack of interest. We compared the responses from individuals who are currently teaching a dedicated bioinformatics course to those who are not. Individuals teaching a dedicated bioinformatics course report that their students lack the appropriate background at a much higher rate than those not teaching a dedicated course (Fig. 5). We hypothesize that faculty at doctoral-granting institutions, who report faculty training to be less of an issue than faculty at other types of institutions, have higher expectations with respect to the skills of their students. On the other hand, lack of interest by students, which we interpret in part to mean students not signing up for courses in bioinformatics among other elective course offerings, may make it difficult to include bioinformatics in the curriculum routinely. To illustrate these barriers—student lack of skills/knowledge and lack of interest—we include relevant quotes below.

- *Student issues: Lack of background skills/knowledge* “The lack of basic mathematical and computational knowledge and skills among students, even in a private university.” “The students’ ability to think critically and effectively evaluate information. They rely on being told exactly what to do and have an inability to problem solve when things don’t turn out as they anticipate.” “Biology [students] shy away from mathematical and statistical methods, and few have programming experience.”
- *Student issues: Lack of interest* “Finding ways to make students interested and engaged in the material presented.” “Student skepticism/resistance.” “Apathy on the part of students.”

**Fig. 5.**
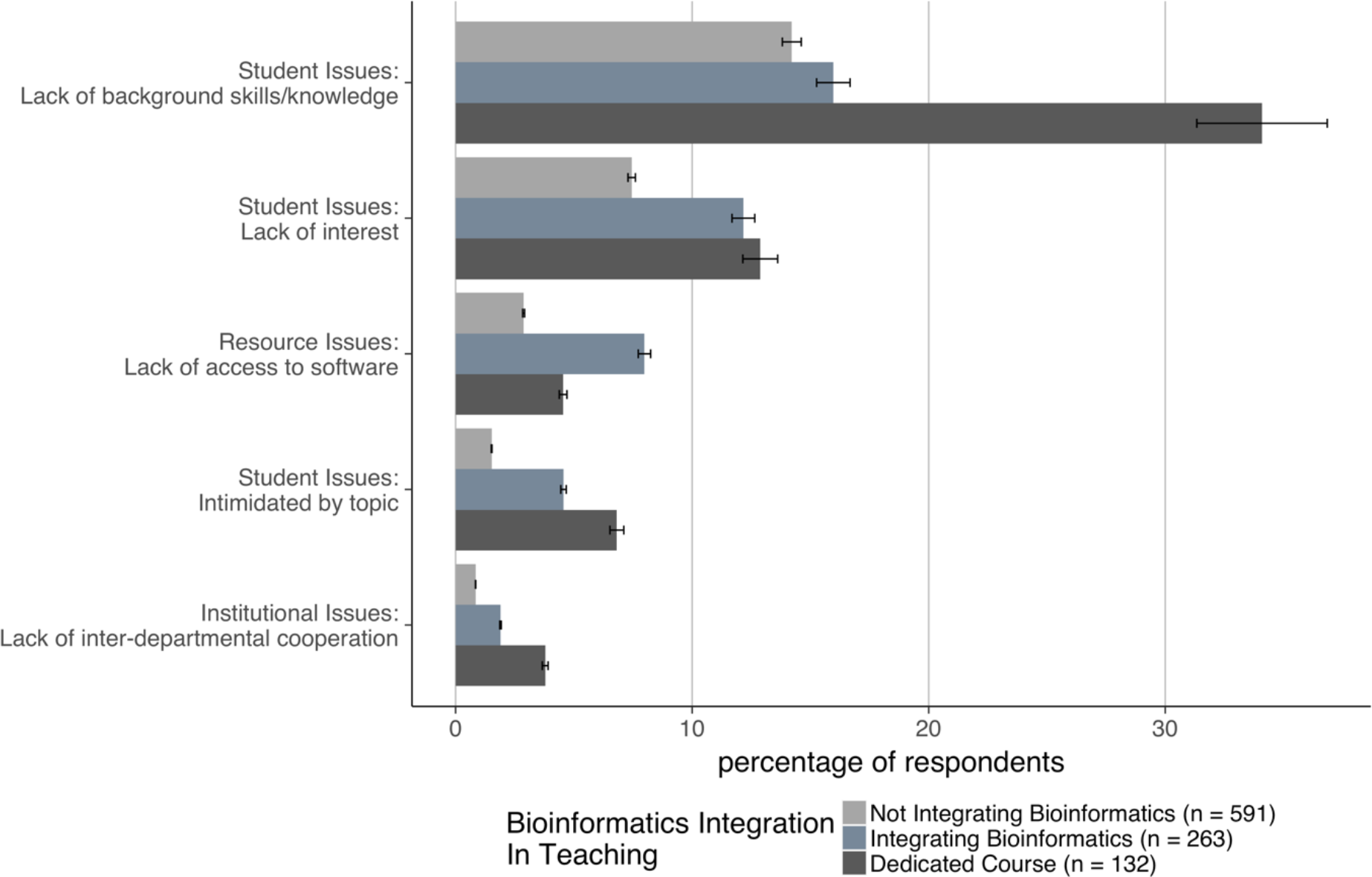
Barriers Reported Grouped by Extent of Bioinformatics Integration into Teaching. Respondents were asked to indicate how (if at all) they were currently integrating bioinformatics into their teaching. Approximately 13% of respondents reported teaching some form of dedicated bioinformatics course; 26% were integrating bioinformatics into their courses in a “substantial” way; and 60% of faculty were not including any “substantial” bioinformatics in their existing courses. Student’s lack of background knowledge and skills (*P* = 2.5e-7) was most frequently reported by faculty teaching a dedicated bioinformatics course (34% vs. 14% of faculty not integrating bioinformatics). Student lack of interest (*P* = 0.03) was reported by at least 10% of faculty (13% of faculty teaching dedicated courses, 12% of faculty integrating bioinformatics). Access to software (*P* = 0.03), student intimidation (*P* = 0.01), and lack of interdepartmental cooperation (*P* = 0.03) were only reported by small numbers of faculty, but differed significantly among cohorts. (*n* = 986, effect size at 80% power = 0.1, meaning small effects were detected).

### Other Barriers

Many faculty also reported issues we grouped under the super-category “Curriculum Issues.” The two categories most frequently mentioned by respondents were “Lack of integration” (cited by about 7% of respondents) and “Too much content” (cited by about 6% of respondents; Fig. 2). Many respondents also mentioned “Quickly changing technologies,” alluding to the difficulties in keeping up with this rapidly changing field, both in terms of training and access to software. This last barrier appeared to be especially problematic for faculty at baccalaureate-granting institutions (Fig. 3), where faculty often have higher teaching loads across a wider range of subjects and have fewer resources. Interestingly, this barrier seemed to be less of an issue at associate’s-granting institutions, which also have high teaching loads, but this may reflect the prescribed curriculum found at two-year institutions.

Finally, several faculty also mentioned issues around institutional support, including fellow faculty who do not feel bioinformatics has a place in the life sciences curricula and lack of support from administrators for resources such as training for faculty or hiring faculty with the appropriate training. The following quote captures these thoughts:

> “Students and other faculty seeing the relevance, understanding that this is where biology is. Letting go of traditional approaches to make room, recognizing that this is an area in which to hire new faculty.”

In Fig. 6, we summarize the data obtained from the respondents, stratified by Carnegie classification. Based on this representation, faculty at associate’s-granting institutions are significantly different from those at the other three types of institutions in a number of aspects. These faculty are least likely to be including bioinformatics in their teaching and more likely to report little to no training in bioinformatics, despite the fact that skills in big-data analysis would contribute to the workforce readiness of their students. In contrast, faculty at doctoral-granting institutions are more likely to have formal training in bioinformatics and to teach dedicated courses in this discipline; they are also the most likely to mention higher-level student issues, such as poor preparation of students in computer science. Faculty at baccalaureate- and master’s-granting institutions are more likely to have obtained training via informal modes of training, including workshops and boot camps. Overall, faculty training is the major barrier preventing integration of bioinformatics instruction into biology curricula for respondents at any type of institution. In the context of training, faculty self-taught in bioinformatics are the most likely group to integrate bioinformatics into their teaching, with 18% of these faculty integrating bioinformatics into existing courses or teaching dedicated bioinformatics courses. Of the faculty identifying workshops and boot camps as their primary bioinformatics training, 13% integrate bioinformatics into their teaching, followed by just 11% of faculty who report some form of formal training. Only a single respondent who indicated no training was incorporating bioinformatics into courses. We conclude that faculty development would be a useful target for future interventions.

**Fig. 6.**
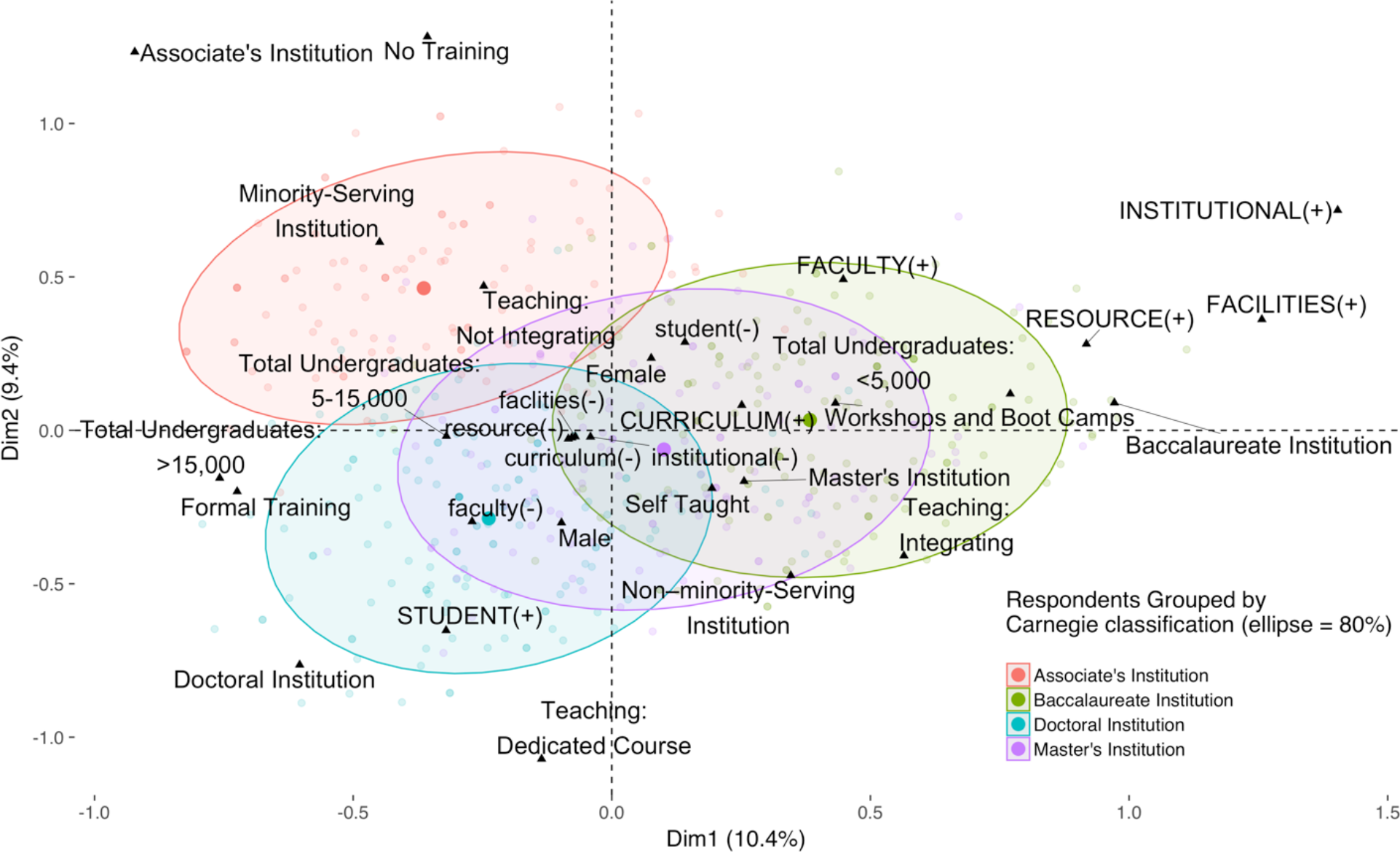
Barriers Reported by Faculty Members Stratified by Carnegie Classification. Multiple correspondence was calculated (see Fig. 4 and Materials and Methods) grouping faculty by Institutional Carnegie Classification (a colored ellipse encompasses 80% of individuals in a sub-demographic, the center of which is marked by a large colored dot). As mentioned in the narrative, the figure shows that faculty at associate’s-granting institutions are different from other institutions in a number of key aspects with respect to barriers to inclusion of bioinformatics in life sciences curriculum. In contrast, faculty at the other institution types map along a continuum, with faculty at baccalaureate-granting institutions more likely to integrate bioinformatics into their teaching, faculty at doctoral-granting institutions more likely to teach dedicated bioinformatics courses, and faculty at master’s-granting institutions in the middle (*n* = 526).

## Discussion

The NIBLSE survey, with more than 1,260 respondents, is the first large-scale survey of life sciences faculty concerning bioinformatics instruction in the United States. In our analysis, faculty training was the most prominent barrier to integrating bioinformatics instruction into undergraduate biology education. This finding was true whether the respondent data were stratified by Carnegie Classification, sex, ethnicity, minority-serving institution status, the size of the undergraduate population, or geographic region. Faculty also often mentioned time, which could be construed in different contexts—time for training, time for instruction, as well as time for restructuring of the curriculum. This latter point could be explored in more detail in future efforts.

In 1998, Altman proposed it was an appropriate time to develop graduate programs in bioinformatics and computational biology (14). While many such programs at the graduate level have been developed since then (15, 16), we see little evidence that graduates from these programs have made an appreciable impact on undergraduate bioinformatics education as faculty to date. Preparing faculty that are equally well-trained in the biology, mathematics, and statistics necessary to teach the breadth of bioinformatics is a long-standing dilemma; initiatives such as QUBES (Quantitative Undergraduate Biology Education and Synthesis) have been addressing this gap (17, 18).

Other studies have also discussed faculty, student, and institutional barriers to integration of bioinformatics into life sciences education. Barone, Williams, and Micklos (19), surveying 704 National Science Foundation investigators from the Directorate for Biological Sciences, also found that training was the top unmet need within the research community. Cummings and Temple (16) describe three general categories of challenges for broader incorporation of bioinformatics in education: (a) required infrastructure and logistics, (b) instructor knowledge of bioinformatics and continuing education, and (c) the breadth of bioinformatics and the diversity of students and educational objectives. Barriers we uncovered here with faculty in the United States are also felt by faculty in the United Kingdom (9), as well as in emerging areas more globally (8), specifically in some African countries (10) and in India (11).

What can be done to alleviate barriers? What can NIBLSE and other such groups do to facilitate solutions? Although there are several institutions that have successfully integrated bioinformatics into their life science programs (20), including University of Wisconsin-La Crosse (21), Kalamazoo College (22), Muhlenberg College (23), and Drake University (24), the vast majority of institutions appear not to have done so. Clearly, given that we and others (16, 25) have found that lack of faculty training is a major problem, providing faculty with opportunities for training is important, as is giving faculty time to take advantage of these opportunities.

At present, there are many opportunities available in the United States and elsewhere, including workshops by groups such as BioQUEST (http://bioquest.org), Data Carpentry (http://www.datacarpentry.org) (26), DNA Subway (http://dnasubway.cyverse.org), Genome Consortium for Active Teaching (GCAT)-Seek (http://gcat-seek.weebly.com) (27), Genomics Education Partnership (http://gep.wustl.edu) (28, 29), Genome Solver (http://genomesolver.org) (30), Integrated Microbial Genomes Annotation Collaboration Toolkit (31, 32), SEA-PHAGES (http://seaphages.org) (33), Software Carpentry (http://software-carpentry.org), QUBES (http://qubeshub.org), the National Center for Biotechnology Information at the National Institutes of Health (http://ncbi.nlm.nih.gov), and the European Bioinformatics Institute (http://www.ebi.ac.uk). Such groups are important not only for conveying information and knowledge but for building community. In addition, many schools offer graduate courses and certificates, either in person or online. There are also numerous courses offered in bioinformatics and computer science through Coursera (http://coursera.org) and EdX (http://edx.org). However, finding these training opportunities is left to individual faculty. NIBLSE plans to serve as a clearinghouse for such opportunities (see the NIBLSE website at https://niblse.org for more information). One of our key findings is that faculty who have participated in informal training like workshops or boot camps report the need for training more than faculty with no training or faculty with formal training. This result is similar to that reported by Feldon et al., who suggest that boot camps and short workshops are not very effective for PhD students in the life sciences (34). Therefore, it may be useful to conduct a follow-up survey to address the deficits expressed by faculty with informal training.

Cummings and Temple (16) recommend “using transformative computer-requiring learning activities, assisting faculty in collecting assessment data on mastery of student learning outcomes, as well as creating more faculty development opportunities that span diverse skill levels, with an emphasis placed on providing resource materials that are kept up-to-date as the field and tools change.” NIBLSE is developing a set of teaching tools that will help contextualize bioinformatics in the light of the fundamentals of biology by developing resources to be stored on the NIBLSE Incubator space at https://niblse.org. We also point to the increasing number of resources in the Bioinformatics course on the *CourseSource* website (http://coursesource.org). These two centers of collected resources will also address the concern exhibited by our respondents as to the difficulty of finding tested curriculum to use in their classrooms. We also note that important fundamental concepts in biology, including evolution and the central dogma, could be taught in the context of bioinformatics, helping to alleviate the “too-full curriculum” barrier.

Cummings and Temple (16) also acknowledge the faculty training issue, saying "Addressing this challenge of instructor knowledge will likely require a multi-fronted approach including: (i) expanding the availability of short (e.g. several days to several weeks) courses focusing on the conceptual background and educational strategies for implementing specific aspects of bioinformatics into courses and curricula; and (ii) the development of lecture material and faculty-customizable web-based student learning activities and associated background materials for deepening faculty understanding.”

Our survey gathered responses from a large number of life science faculty, but we note there were few respondents who identified as members of groups underrepresented in STEM; despite this, the numbers are consistent with percentages of such faculty members at institutions around the country (13). Although we are aware that members of individual groups may have different needs, responses from underrepresented groups were binned together because the numbers of respondents from these groups was low (81 total respondents). Previous reports have noted that at many historically black colleges and universities, bioinformatics courses have not been implemented widely due to a number of factors similar to those outlined here for the wider range of faculty, including lack of faculty training and lack of resources (35).

The NIBLSE survey revealed a number of issues that life sciences faculty identified as barriers to integration of bioinformatics into the curriculum. A future survey could take advantage of these findings by posing specific questions about the extent to which individual faculty experience specific barriers. Faculty training in bioinformatics is the major barrier reported by faculty, but given that more graduate students are receiving training in this area currently, we predict this barrier should eventually resolve itself as more of these students move into faculty positions. However, since the youngest respondents report teaching bioinformatics the least, perhaps these new faculty need to acquire enough seniority to update the curriculum. Thus, we believe that as these faculty members mature, bioinformatics will become a standard part of the life science curriculum. Until then, NIBLSE is committed to working to eliminate barriers by making faculty aware of training opportunities that currently exist, potentially developing additional training opportunities, and identifying and curating resources for integrating bioinformatics into life sciences education more broadly.

In sum, the results of this survey of more than 1,260 life science faculty show the vast majority agree that bioinformatics should be integrated into the undergraduate life sciences curriculum. However, our results further indicate lack of faculty training is the single biggest barrier to integration. Surprisingly, the faculty with the most training in this area—those in the cohort of faculty who received their PhD degrees most recently—are teaching bioinformatics the least. In addition, the survey results reveal a significant gap in the landscape of bioinformatics education and indicate that there is a critical need for quality faculty training. This need is especially great for faculty who are members of underrepresented groups in STEM and for faculty at associate’s-granting institutions. While many questions remain, moving forward, NIBLSE seeks to address the challenges uncovered in the present analysis in order to achieve integration of bioinformatics into the life sciences curriculum. We invite those with interest in this issue to join us (http://nibles.org).

## Materials and Methods

The survey of life science faculty was collaboratively developed by a sub-group of NIBLSE members, the Core Competencies Working Group (CCWG). Faculty from a range of educational institutions were represented in the CCWG, including faculty at baccalaureate-granting, master’s-granting, and doctoral-granting institutions with various levels of research activity. One of the members of the CCWG was from industry. All members of the committee have extensive experience teaching bioinformatics to undergraduate life science students. Development and deployment of the survey is discussed in more detail in our companion paper (submitted, accessible at http://www.biorxiv.org/content/early/2017/08/03/170993; the survey in its entirety is provided as a Supplementary Document in this paper). Approval for the study was obtained from the University of Nebraska at Omaha Institutional Review Board (IRB # 161-16-EX) before the survey was distributed.

The survey was administered in April 2016 using Qualtrics with assistance from the Center for New Designs in Learning and Scholarship at Georgetown University; 1,264 responses were collected. The branched survey design included five-point Likert and free-response questions. As described in detail in our companion paper, the survey was sent to more than 11,000 email addresses of biology faculty purchased from MDR (http://schooldata.com) and to members of networks of faculty with interests in life sciences education. Using information provided to us by MDR and from publicly available databases (36-38), we estimate that there are currently between 75,000 and 100,000 biological sciences faculty in the United States. Given our overall sample size (1% to 2%) we estimate that the mean margin of error for the survey questions described in this paper is ±3% at the 95% confidence interval (Fig. 1) (39). For the results described here, we analyzed barriers to teaching bioinformatics through four free-response questions (Table 1). These responses were subjected to qualitative analysis by two groups, one at Georgetown University (AGR, using methods outlined in Leech and Onweugbuzie [40], specifically the classic content analysis) and one at the University of Florida (JCD, SG, and EWT, using a modification of the coding and thematic analysis process described by Harding [41]). In both analyses, keywords were identified and organized into categories (e.g., “no expertise/training,” “time,” “not enough faculty”) and combined into super-categories (e.g., “Faculty Issues,” “Student Issues,” “Curriculum Issues,” “Resource Issues,” “Facilities Issues,” “Institutional/Departmental Support Issues”) as shown in Table S1 for Question 1. The number of responses were scored relative to the keywords. Although similar results were obtained from the two analyses, the authors agreed to use the data from the University of Florida quantification for detailed analyses because the way in which the data was formatted made subsequent analyses easier.

Survey data were exported to CSV-formatted files for analysis in R. Data were cleaned to eliminate multiple column headers and to transform Qualtrics numerical coding of responses into decoded values. During this step, responses from outside of the United States were eliminated (leaving 1,231 valid responses). In some cases (i.e., respondent race/ethnicity, level of bioinformatics training, and degree year), responses were binned to achieve sufficient numbers for analysis (e.g., all survey respondents from races/ethnicities underrepresented in STEM were analyzed together).

**Analysis:** The reported barriers for each question were analyzed with respect to a number of demographic criteria (i.e., sex, race/ethnicity, highest degree earned, year of highest degree, level of bioinformatics training, extent of current bioinformatics teaching, institutional Carnegie classification, minority-serving institution [MSI] vs. non-MSI status, size of school by undergraduate enrollment, and geographic region) to determine differences within these demographics and association of demographics and barriers. For a given demographic, respondents who did not answer, or indicated they did not know or were unsure, were dropped from analysis of that demographic category.

Multiple correspondence analysis (MCA) was used to visualize the correspondence of several categorical demographic factors (42). Similar to a principle component analysis, MCA allows associations between qualitative variables (e.g., our demographic categories) to be visualized. In our analysis, individuals for which we had complete demographic data were used display relationships in two-dimensional space.

**Proportion tests within demographics:** A proportion test was used to calculate the Chi-squared statistic for differences between sub-demographics (H_0_ assuming that faculty within all the sub-demographics report barriers equally). The margin of error (as the interval estimate of population proportion) was calculated at the 95% confidence level and is represented on the figures (Figs. 3, 5, and S3) as error bars. Expected effect sizes detectable were calculated assuming 80% power. Selected findings are described in Results; additional findings as well as the full data set and R scripts used for analyses and plotting can be found on the NIBLSE GitHub repository (43).

## Acknowledgements

This material is based upon work supported by the National Science Foundation under Grant Number 1539900 to AGR, EWT, ED, WM, and MAP. Any opinions, findings, and conclusions or recommendations expressed in this material are those of the author(s) and do not necessarily reflect the views of the National Science Foundation. The authors thank the members of the Genomics Education Partnership, Genome Solver, GCAT-SEEK, and NIBLSE networks for the feedback they provided. AGR thanks Gopal Topiwala for his help with the Georgetown analysis. JCD, SG, and EWT thank Jonathan Orsini for his help with the UF analysis. We also thank Courtney Soderberg and the statistical consulting service at The Center for Open Science.

